# A rapid platform for 3D patient-derived cutaneous neurofibroma organoid establishment and screening

**DOI:** 10.1101/2022.11.07.515469

**Authors:** Huyen Thi Lam Nguyen, Emily Kohl, Jessica Bade, Stefan E. Eng, Anela Tosevska, Ahmad Al Shihabi, Jenny J. Hong, Sarah Dry, Paul C. Boutros, Andre Panossian, Sara Gosline, Alice Soragni

**Affiliations:** Department of Orthopaedic Surgery, David Geffen School of Medicine, University of California Los Angeles, CA, USA; Pacific Northwest National Laboratories, Seattle, WA, USA; Department of Human Genetics, University of California Los Angeles, CA, USA; Department of Molecular, Cell and Developmental Biology, University of California Los Angeles, CA, USA; Department of Pathology, David Geffen School of Medicine, University of California Los Angeles, Los Angeles, CA, USA; Division of Hematology-Oncology, David Geffen School of Medicine, University of California Los Angeles, CA, USA; Institute for Precision Health, University of California Los Angeles, CA, USA; Department of Urology, University of California Los Angeles, CA, USA; Jonsson Comprehensive Cancer Center, University of California Los Angeles, Los Angeles, CA, USA; Eli and Edythe Broad Center of Regenerative Medicine and Stem Cell Research, University of California Los Angeles, CA; Panossian Plastic Surgery, Pasadena, CA, USA; California NanoSystems Institute, University of California Los Angeles, CA, USA; Molecular Biology Institute, University of California, Los Angeles, Los Angeles, CA, USA

**Author notes:** Correspondence to SG and AS.

## Abstract

Localized cutaneous neurofibromas (cNFs) are benign tumors that arise in the dermis of patients affected by Neurofibromatosis Type 1 syndrome (NF1). cNFs are fundamentally benign lesions: they do not undergo malignant transformation or metastasize. Nevertheless, in NF1 patients, they can cover a significant proportion of the body, with some individuals developing hundreds to thousands of lesions. cNFs can cause pain, itching, and disfigurement with substantial socio-emotional repercussions. To date, surgical removal or laser desiccation are the only treatment options, but can result in scarring and the leave a potential for regrowth.

To support drug discovery efforts focused on identifying effective systemic therapies for cNF, we introduce an approach to routinely establish and screen cNF tumor organoids. We optimized conditions to support ex vivo growth of genomically-diverse cNFs. Patient-derived cNF organoids closely recapitulate the molecular and cellular heterogeneity of these tumors as measured by immunohistopathology, DNA methylation, RNA-seq and flow cytometry. Our tractable patient-derived cNF organoid platform enables rapid screening of hundreds of compounds in a patient- and tumor-specific manner.

## Introduction

Neurofibromatosis Type 1 is an autosomal dominant hereditary syndrome caused by germline mutations in the neurofibromin gene^1–3^ (*NF1*). *NF1* is a tumor suppressor gene located on chromosome 17q11.2, where it occupies over 280kb across 61 exons. The gene encodes a 2,818 amino acid GTPase activating protein and negative regulator of RAS^4^. Germline variants as well as sporadic acquired somatic mutations in the *NF1* gene can lead to elevated levels of RAS-GTP levels, increased RAS signaling and uncontrolled cellular proliferation^5,6^.

Deleterious germline variants in this tumor suppressor affect 1:2,500 to 1:3,500 individuals, leading to an array of symptoms including pain, cognitive issues, and the growth of benign tumors throughout the body with different potential for malignant transformation. Over 90% of individuals with Neurofibromatosis Type 1 develop localized cutaneous neurofibromas (cNFs), which are benign cutaneous growths with no risk of progression to malignant, invasive disease^7,8^. cNFs typically emerge in the second half of the first decade of life, grow or expand in numbers during puberty and can number in the thousands^7–9^. While local invasion or development of metastases are never observed for this type of benign tumors, they can give rise to pain, itching, and disfigurement, directly impacting the quality of life of patients^10^. Importantly, there is no systemic treatment option available to NF1 patients for treating cNF lesions^8^. Interestingly, not all patients are equally affected by cNFs^10–12^, which is linked, in part, to the high degree of genetic diversity in the *NF1* alterations reported^12,13^. The most common mutation in *NF1*, p.Arg1809Cys, is found in only ∼1% of unrelated individuals in a ∼7,000 patient cohort^12,13^. Overall, patients with microdeletions have higher tumor volumes while patients with p.Arg1809 codon alterations tend to have fewer lesions and those with p.Met992del have no cNFs at all^12,13^. Additional variability can arise from timing and type of secondary alterations, hormonal factors (cNF arise during puberty and can worsen in pregnancy), and contributions of the microenvironment^14,15^. As such, there are several outstanding questions surrounding cNF origin, manifestation and development^14^.

The mutational heterogeneity has led to difficulties in developing clinically relevant model systems. A transgenic *Sox10*-CreERT2 *Nf1*^fl/fl^ mouse model that can develop cNFs was recently established^16^. A mouse model with targeted *NF1* knockout in *Prss56*-expressing boundary cap cells located in the neural crest, gives rise to cNF-like lesions in adult mice^17^. In addition, two porcine models have been reported, including a heterozygous *NF1*^R1947^ mutant and a *NF1*^+/ex42del^ carrying a deletion of exon 42^18,19^. Despite these recent advances, *in vivo* models lack the ability to fully recapitulate the characteristics and genomic diversity of human cNFs and are impractical for large scale drug screening.

In addition to the genetic heterogeneity, cNFs are a mixture of diverse cell types including Schwann cells, fibroblasts, mast cells as well as pericytes and endothelial cells^14,20^. A recent single-cell RNAseq analysis has confirmed the cellular heterogeneity of cNFs and the role of specific collagens in supporting cNF growth^21^. As such, any *ex vivo* or *in vitro* cNF model must recapitulate such cellular diversity and extracellular matrix components, should accommodate different *NF1* alterations and be amenable to high-throughput screening.

Tumor organoids are exquisitely suited for modeling heterogeneous tumors; these are tractable models of disease generated from patient material that can maintain genome alterations and faithfully recapitulate the histopathology of the parent tissue^22^. We developed a platform for the rapid establishment of tumor organoids from both aggressively growing^23^ and indolent tumors^24^. By taking advantage of a modified geometry, organoids can be established and grown in rings of extracellular matrix around the rim of wells, a design pattern compatible with automation and high-throughput drug screening protocols^23–25^. Here, we leverage this platform to grow and screen cNF organoids. Firstly, we determine the conditions that support *ex vivo* growth of diverse cNFs while best recapitulating the parental tumor in terms of cellular heterogeneity, transcriptome, methylome, and protein expression profile. Second, we perform a proof-of-principle screening to confirm feasibility to implement the mini-ring screening platform and identify pathways susceptible to inhibition in benign lesions. Overall, we show how this technology can be applied for drug discovery studies and therapeutic development targeting benign, diverse tumors such as cNFs.

## Results

### cNF organoids can be established and recapitulate the immunohistopathology of the tumor of origin

We enrolled n=12 NF1 patients undergoing cNF removal as part of clinical management. Of these, 8/12 (66%) were female, and median age was 52 (range 29-65, **Table 1**). This distribution is in line with the typical characteristics of patients who are affected by cNFs. cNFs were collected from different body locations (**Table 1**), and ranging in size, stiffness, and appearance (**Figure 1**). Overall, we processed over 100 cNFs; those with sizes greater than 0.5 cm were processed individually and yielded between 0.3 – 7.5 million cells/ cNF. Smaller tumors were pooled, with multiple cNFs from the same patient processed together. In all cases, a portion of the cNF is preserved for immunohistochemistry and sequencing studies. The remaining portion of cNFs were dissociated to single cells and reconstituted *ex vivo* (**Figure b**). Several samples yielded sufficient cells to seed organoids and perform optimization studies as well as drug screenings using our established approach^23–25^ (**Figure 1b**). Overall, 100% of the samples tested grew well in our ring organoid system, giving rise to viable organoids within a week ofseeding (**Figure 1c**).

**Table 1.**
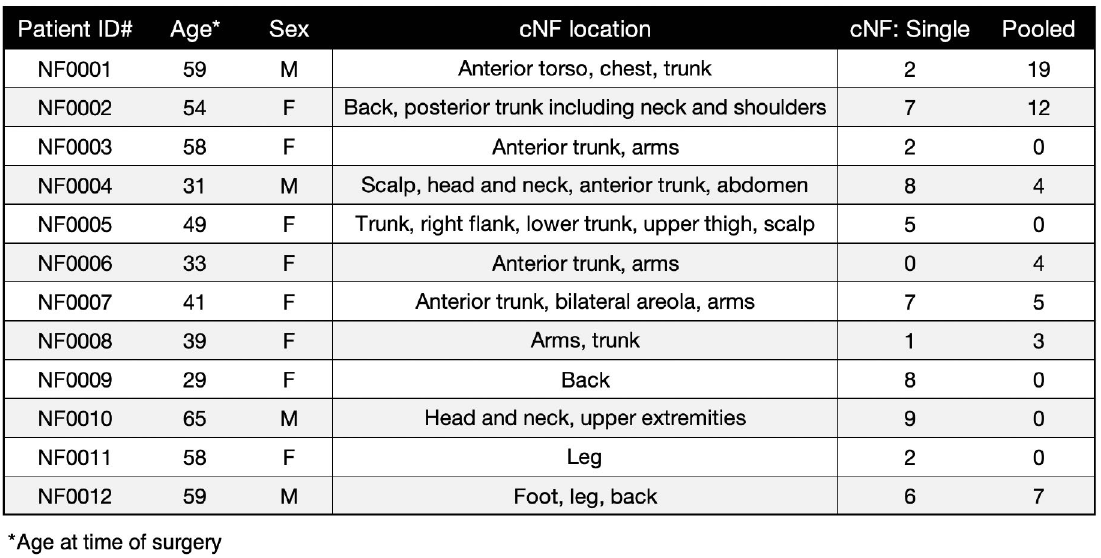
Characteristics of enrolled patients and corresponding cNFs.

**Figure 1.**
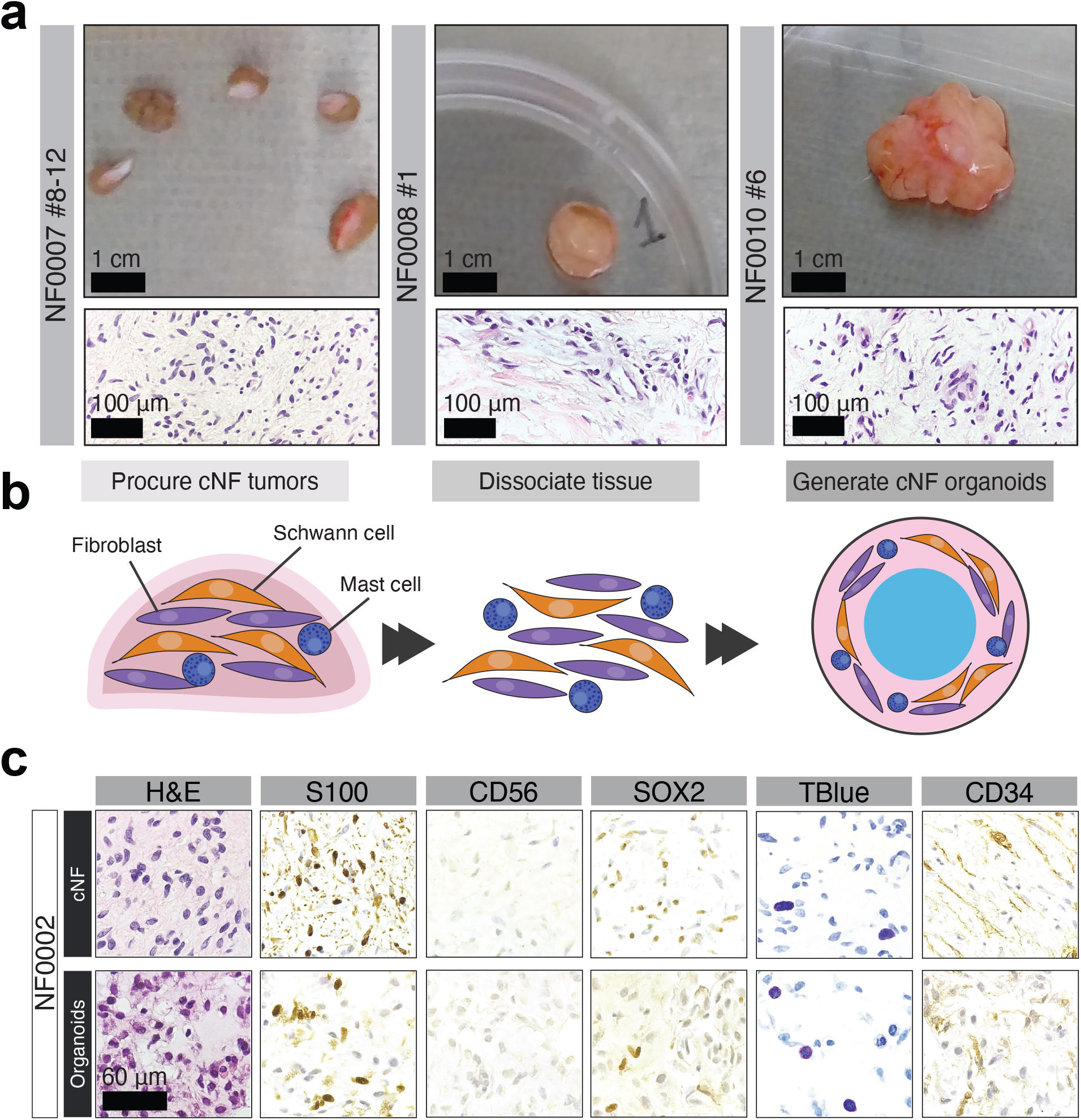
Clinically diverse cNFs give rise to organoids with similar histopathological features. **(a)** cNFs have diverse sizes and stiffness, appearance on H&E staining. **(b)** Schematic of the process to dissociate cNFs and reconstitute them ex vivo in ring format for facile screening. **(c)** Immunohistopathology of a cNF from patient NF0002 and the corresponding organoids shows the heterogeneous cell populations are retained upon culturing in our system. S100 stains Schwann cells, CD56 is a neuronal marker, SOX2 is a marker for dedifferentiated Schwann cells, Toluidine Blue is indicative of mast cells and CD34 of fibroblasts.

cNFs are composed of many different cell types, the most abundant being Schwann cells, fibroblasts, and mast cells^14,20^. Immunohistopathology of the cNF organoids shows that the cell populations within the organoids closely match the cells in the cNFs of origin (S100+, CD56-, CD34+, SOX2+, Toluidine blue+, **Figure 1c**). Expression of these markers is typical of the histopathology of cNF tumors^14,26^. For instance, we observed extensive S100 positivity, indicative of Schwann cells, a lattice-like CD34+ fibroblastic network^27^, and mast cells as visualized by toluidine blue staining^28^ (**Figure 1c**).

#### cNF organoids can be established regardless of genomic variant

We selected five patients with sufficient available cells for a thorough growth condition optimization screening and downstream characterization. First, we performed whole genome sequencing to determine the *NF1* variants present as well as any pathogenic variant in other genes (**Table 2, Figure 2a**).

**Table 2.** Summary of pathogenic variants found by WGS.

| Patient ID# | Gene | Variants identified | Location |
| --- | --- | --- | --- |
| NF0002 | NF1 | Large chromosomal deletion | chr17:31234616-31238587 |
| NF0007 | NF1 | Structural variant | chr17: 31332573-31332924 |
| NF0007 | NF1 | Splicing variant SNV | chr17: 31163377 G->A |
| NF0007 | DYSF | Stop gain SNV | chr2: 71570679 C->T |
| NF0007 | TNFRSF13B | Stop gain SNV | chr17: 16948752 G->C |
| NF0008 | NF1 | Splicing variant SNV | chr17:31330295 G->T |
| NF0008 | COL6A3 | Splicing variant SNV | chr2: 237387581 C->T |
| NF0008 | GAA | Non-synonymous SNV | chr17:80112604 G->A |
| NF0009 | NF1 | Structural variant | chr17: 31332573-31332924 |
| NF0009 | SOS1 | Splicing variant SNV | chr2: 38989315 C->T |
| NF0012 | NF1 | Structural variant | chr17: 31332573-31332924 |
| NF0012 | NF1 | Stop gain SNV | chr17: 31206360 C->T |
| NF0012 | BBS10 | Nonsynonymous SNV | chr12: 76348214 G->A |

**Figure 2.**
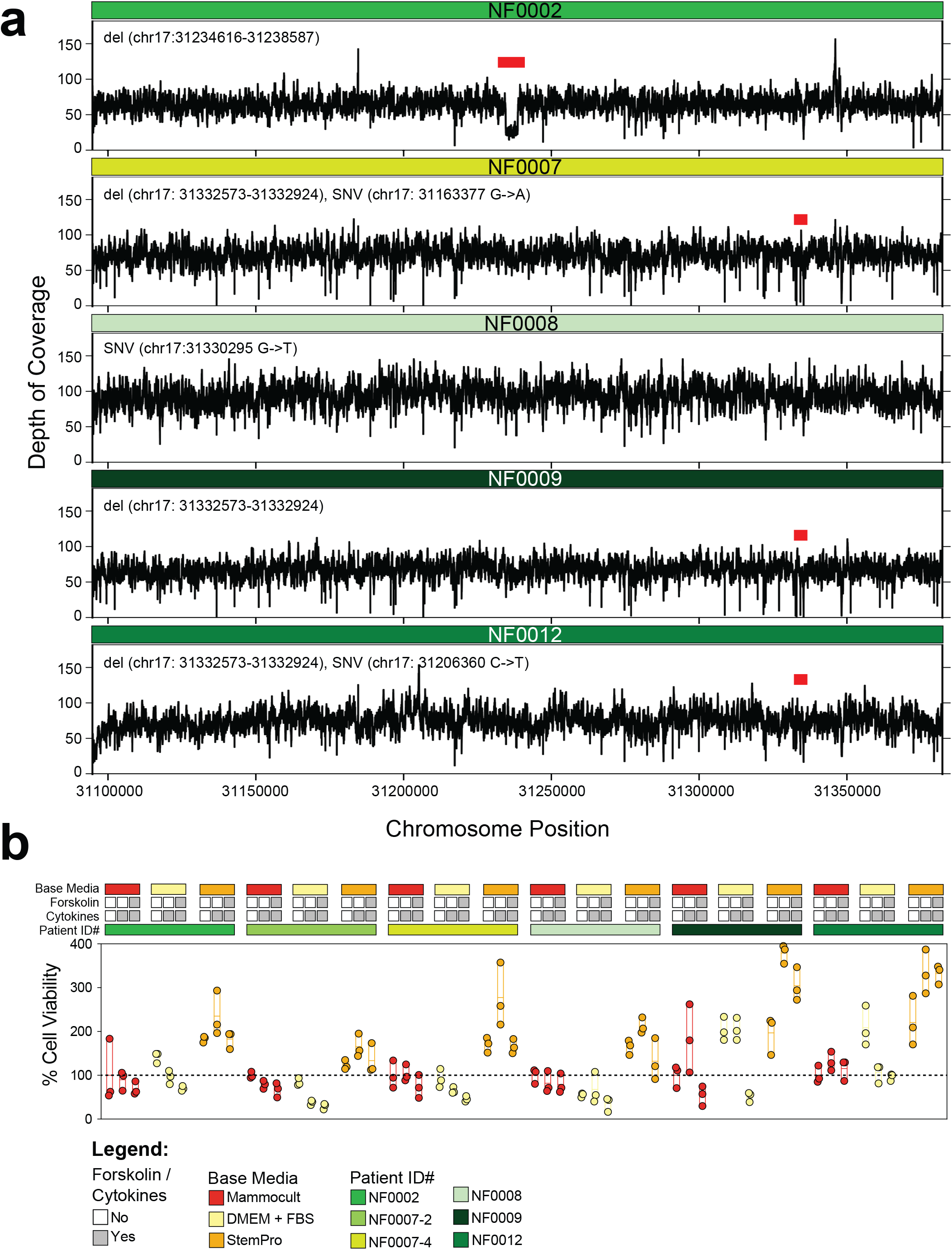
cNFs harboring NF1 alterations generate viable organoids. **(a)** Coverage of the *NF1* gene as estimated by GATK4 is shown for each subject. **(b)** All samples generated viable organoids in culture as determined by end-point ATP-release assay, regardless of the NF1 alteration present. Different media sustain the growth of cNF organoids, with StemPro-based formulations promoting strong proliferation.

Four out of the five patients sequenced exhibited pathogenic structural alterations in the *NF1* gene. NF0002 has a substantial dip in coverage between position chr17:31234559-31237611, due to a large deletion (**Figure 2a, red bar**). Patients NF0007 and NF0012 each had two identifiable *NF1* alterations, a shared deletion (chr17: 31332573- 31332924) coupled to a splicing variant (NF0007) or a stop gain single nucleotide variant (NF0012). Interestingly, NF0008 and NF0009 each had a single identifiable alteration in *NF1*. While we cannot distinguish between germline and somatic variants, it is likely that structural variants in the germline could impede the ability to identify the second hit SNVs in the NF1 gene. Alternatively, it is possible pathogenic variants in other genes (GAA in NF008, SOS1 in NF0009) may contribute to some extent to the phenotype. Regardless of genotype, we could successfully establish sufficient viable organoids to perform a media optimization screening.

#### Culture conditions affect cNF organoid growth

Given the cellular heterogeneity that is integral to the biology of benign cNFs, we elected to identify conditions that would support the growth and maintain the proportion of each cell type in 3D organoid format. For this, we established nine conditions in total to screen on n=6 samples from five patients. The selection included three base media, StemPro, DMEM with 10% FBS or Mammocult, either alone or supplemented with a cytokine cocktail or cytokines and forskolin. These conditions were selected on the basis of previous work for their ability to favor the growth of cell types similar to those found in cNFs (**Figure S1**).

We first tested if the different media had any differential effect on cell growth and proliferation. Following our platform approach and timeline^23–25^, we seeded cells to develop organoids in 96 well mini-ring format^25^, incubated them for six days in the nine media conditions and measured aggregate viability as endpoint assay (**Figure 2b**). Different culturing conditions and cytokine supplementation significantly affected organoid growth rates (**Figure 2b**). cNF organoids in Mammocult and DMEM exhibited less robust growth patterns in the presence of supplements. On the contrary, a distinct increase in proliferation can be observed when cells were cultured in StemPro, particularly with cytokines and/or forskolin (**Figure 2b**). Thus, StemPro-based media are supporting robust growth of cNF organoids *ex vivo*.

#### Molecular characterization of cNF organoids

Although viability results indicate that StemPro strongly favors cNF cell growth (**Figure 2**), there is a possibility that cells may be selected for in a manner that does not recapitulate the biology of these tumors. As such, we set to quantify the overall transcriptional changes that occurred in all growth conditions and compare these to the cNF of origin. We determined gene expression by performing RNA-seq on 100,000 cells either freshly extracted from a cNF or after establishing organoids and growing them in the different conditions for six days. We then benchmarked each measurement to that from the same primary tumor using Spearman rank correlation statistics. This allows to separate out correlation measurements by culture condition and determine which medium leads to the best concordance between organoid models and patient samples. Our results show that, on average, cNF organoids correlate strongly with the parental tumors in all conditions tested, with calculated Spearman’s rank correlations > 0.8 (**Figure 3a**). There is a trend toward higher correlation values between parental cNF with cNF organoids grown in StemPro for all samples tested, with a median of 0.91 compared to 0.88 and 0.87 for Mammocult and DMEM respectively, though the difference is not statistically significant (orange dots, **Figure 3a**). The addition of cytokines and/or forskolin had negligent effects (**Figure 3a**).

**Figure 3.**
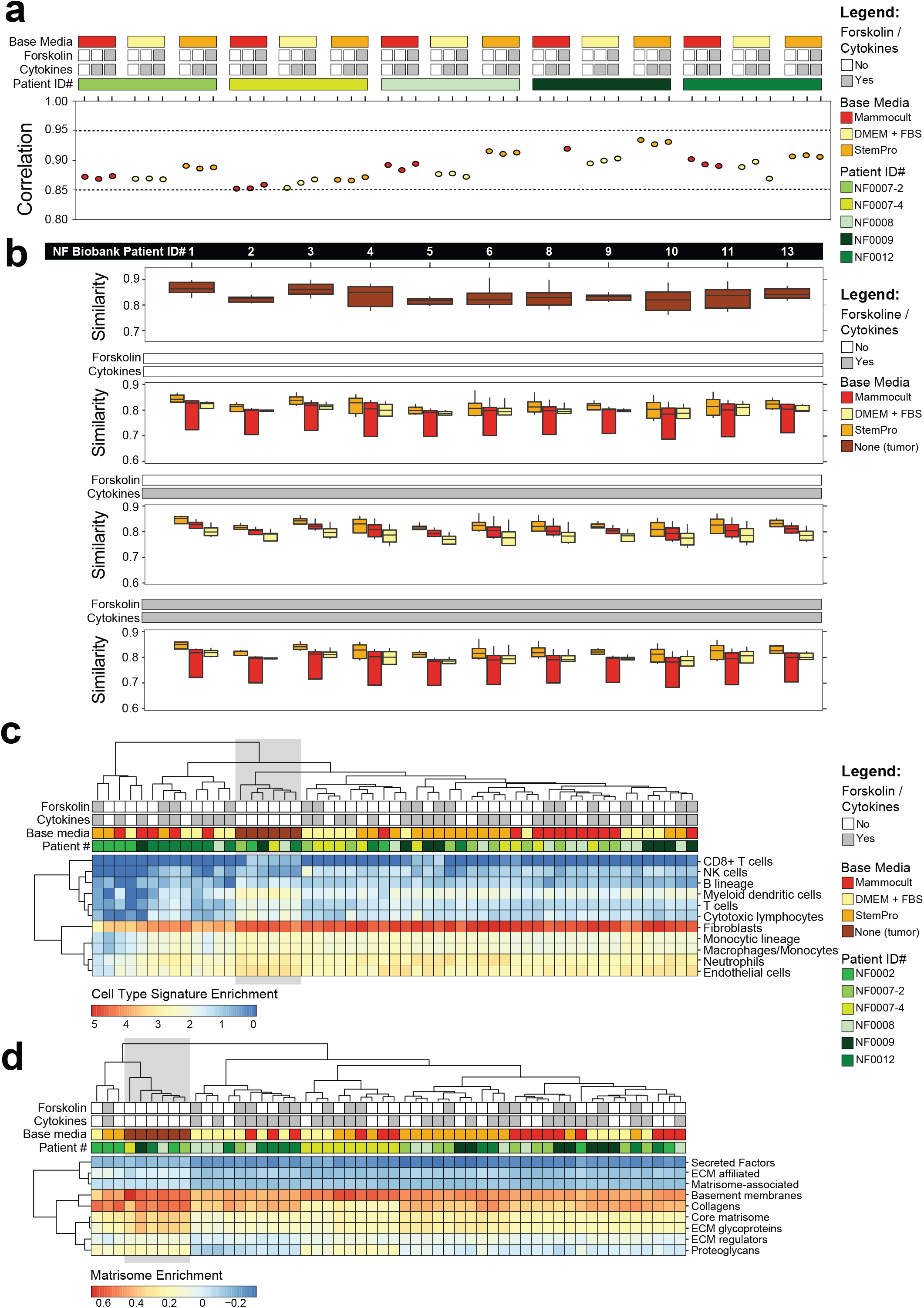
Gene expression analysis and cell type deconvolution. **(a)** Spearman rank correlation of each organoid with the primary tumor for that patient sample. Legend depicting sample properties is on the right. **(b)** Correlation of all tumors (3-4 per patient) from the NF biobank with primary tumor (top row) and organoids (bottom 3 rows). Color indicates base media, bars across top indicate additions to matrix. **(c)** Heatmap of MCPCounter scores representing relative proportion of estimated cell types in each sample. Color legend below. **(d)** Normalized enrichment scores of each sample for the matrisome gene lists listed along the right.

As independent validation of our cohort, we benchmarked gene expression measurements to an existing repository of cNF transcriptional profiles. Specifically, we leveraged the CTF cNF Biobank^15^ which contains n=33 RNA-seq profiles from 11 patients and is publicly available on the NF Data Portal^29^. Our results show that all primary cNFs are highly similar to the historical cohort, with Spearman rank correlations above 0.8 (**Figure 3b, top panel**). Next, we averaged all patient samples grown in the same condition and assessed how well each medium compared to tumors from the CTF cNF repository (**Figure 3b, bottom panels**). Overall, organoid models correlate with most CTF tumors with Spearman’s rank correlation of > 0.6. Yet again, StemPro exhibited the highest correlation values at any supplementation level, with a median correlation of 0.82 compared to 0.8 for both Mammocult and DMEM (**Figure 3b, bottom panels**).

In addition to the transcriptome analysis, we also quantified and compared cNF samples at the methylome level (**Supplementary Table 1**). Changes in methylation across the genome are a sensitive indicator of the effect of growth conditions on cells, and have been observed for long time cultures and repeat passaging^30,31^. Although our platform entails growing the cNFs for only 6 days with no passaging, global loss in 5-hydroxymethylcytosine (5hmC) has been reported within days in culture^32^. Thus, we set to perform Reduced Representation Bisulfite Sequencing (RRBS)^33^ to identify a set of methylated CpG islands across the genome for each sample. We compiled a list of all CpG sites with at least 10x coverage across both the organoids and the primary tumors. We then computed the Pearson correlation of the methylation quantities between the cNF organoids grown in each condition and the corresponding paternal tissue (**Supplementary Table 1**). All media displayed similar correlation values across the overlapping set of CpG islands. The correlation followed the same trend as above, with StemPro having the higher median value (0.72) when compared to DMEM (0.69) or Mammocult (0.71).

### Cell composition analysis confirms that cNF organoids contain multiple cell populations

Our IHC results are indicative of the presence of populations of Schwann cells, fibroblasts, and mast cells in the cNF organoids (**Figure 1c**). We set to provide a quantitative assessment of the amount of each cell type within the cNF lesions and organoids. This is particularly important considering the data on different tumor types showing how sub-populations of cells grown *ex vivo* can rapidly evolve or be selected for as a function of media conditions^34^. Toward this end, we have quantified the percentages of Schwann cells, fibroblast and mast cells with flow cytometry by simultaneously sorting cNFs and derived organoids with 3 markers: S100 for Schwann cells^35,36^, CD34 for fibroblasts^36^ and c-Kit for mast cells^37^ (**Supplementary Figure 2**). There are variable proportions of cells in the parental cNFs, with dominant proportions of S100+ and CD34+, and a smaller amount of c-Kit+ cells, accounting for less than 10%. In cultured organoids, we observe a trend towards a reduction in the amount of CD34+ cells, and occasionally an increase in c-Kit+ cells in some conditions, with the exclusion of StemPro culture conditions.

Next, we leveraged RNA-seq data to compare cell type compositions within each sample by applying a tumor deconvolution algorithm, the Microenvironment Cell Populations-counter (MCP-Counter)^38^. This method allows to identify stromal cell subtypes in bulk transcriptome data on the basis of existing, well characterized refer(ence profiles^38^. While a Schwann-like cell profile is not available, the method could identify the presence of abundant fibroblasts as well as cells from the myeloid lineage, which includes mast cells, in all parental cNFs and most corresponding organoids (**Figure 3c**). This is largely recapitulating our experimental evidence based on immunohistochemistry (**Figure 1c**) and flow cytometry (**Supplementary Figure 2**). Organoids from NF0012 and few media conditions for the ones established from NF0002 had a reduced proportion of myeloid cells/mast cells (**Figure 3c**). In addition to fibroblasts, endothelial cells are also prominent (**Figure 3c**), which is in line with the biology of cNF^14,20^. Proportions of these cells are maintained within the majority of cNF organoids tested. Interestingly, MCP-Counter identified cells compatible with macrophages and neutrophil signatures. Macrophages have been shown to have a role in neurofibroma and are found amongst the inflammatory immuno-infiltrates in cNFs^39,40^. Neutrophils have also been reported in cNF^41^.

Lastly, we interrogated the abundance of extracellular matrix protein transcripts, i.e. the matrisome^21^. Over 50% of the mass of a typical cNF is composed of secreted extracellular proteins, particularly collagen VI which is secreted by resident fibroblasts although all cell types present in cNFs were found to contribute to secretion of the matrisome^21^. To characterize the RNAs encoding extra-cellular matrix proteins, we scored each sample for enrichment of genes encoding for matrisome proteins^42^ using the Gene Set Enrichment Analysis (GSEA) algorithm^43^. The analysis confirms that the core extracellular proteins and collagen genes are generally expressed at similar levels within cNFs and derived organoids (**Figure 3d**).

#### Molecular and growth-based analyses identify the optimal cNF growth conditions

Standardized media growth and screening conditions that can be broadly applied to tumors from different individuals and genomic backgrounds facilitate drug discovery efforts^23,24,44^. As such, we set to determine the media that, in aggregate, are best recapitulating the molecular features of the parental cNFs by combining our phenotypical and molecular analyses (**Figure 2-3 and Supplementary Figure 2**). To do so, we leveraged the Spearman’s rank correlation statistic and compared each organoid measurement (RNA-Seq, Flow Cytometry, Methylation) to the primary tumor (**Figure 4**).

Overall, RNA-Seq showed the highest levels of correlation (median correlation of 0.91) although flow cytometry (0.50) and RRBS (0.72) followed a similar trend (**Figure 4**). Results show how StemPro-based conditions generally gave rise to high correlation values for the largest number of patient-derived cNF organoids samples. While the high correlation values were not statistically significant for each individual dataset, when we pooled the measurements across data types, the correlation values for StemPro were statistically significant in a linear model (p=0.04). This, together we the measured ability to support a robust *ex vivo* cNF organoid growth and proliferation (**Figure 2b**), led us to select StemPro with cytokines +/- forskolin as the media of choice for drug screenings.

**Figure 4.**
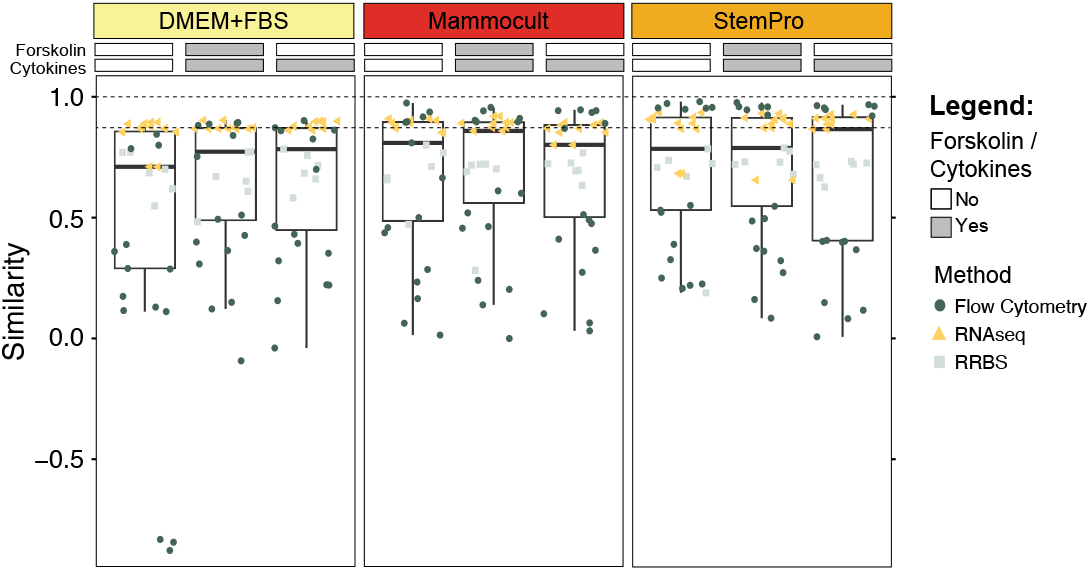
Overall correlation of cNF organoids with parental tissue by data modality and base medium. Each point represents the correlation of an organoid sample with the respective cNF lesion. The color and shape of the points indicate the different data stream used to calculate the correlation.

### High-throughput drug sensitivity screening

Given the established growth conditions, we determine feasibility to perform medium- to high-throughput drug screening on cNF benign tumor organoids. Using our established organoid screening paradigm and mini-ring platform^23–25,45^, we tested n=43 targeted kinase inhibitors on cNF organoids established from two patients (**Figure 5**). NF0009 were tested in StemPro with cytokines while NF0012 was tested in StemPro with cytokines and forskolin. Overall, cNF organoids could be screened with a protocol akin to that used for malignant tumors^23–25^. Both organoid models were sensitive to our positive control, the pan-kinase inhibitor staurosporine, as well as copanlisib (PI3K inhibitor) and onalespid (Hsp90 inhibitor, **Figure 5**). NF0009 showed unique sensitivity to dasatinib (Abl, Src and c-Kit inhibitor) and linsitinib (IGF-1R inhibitor), among others (**Figure 5**). Overall, our approach can be used to determine shared features as well as individual sensitivities.

**Figure 5.**
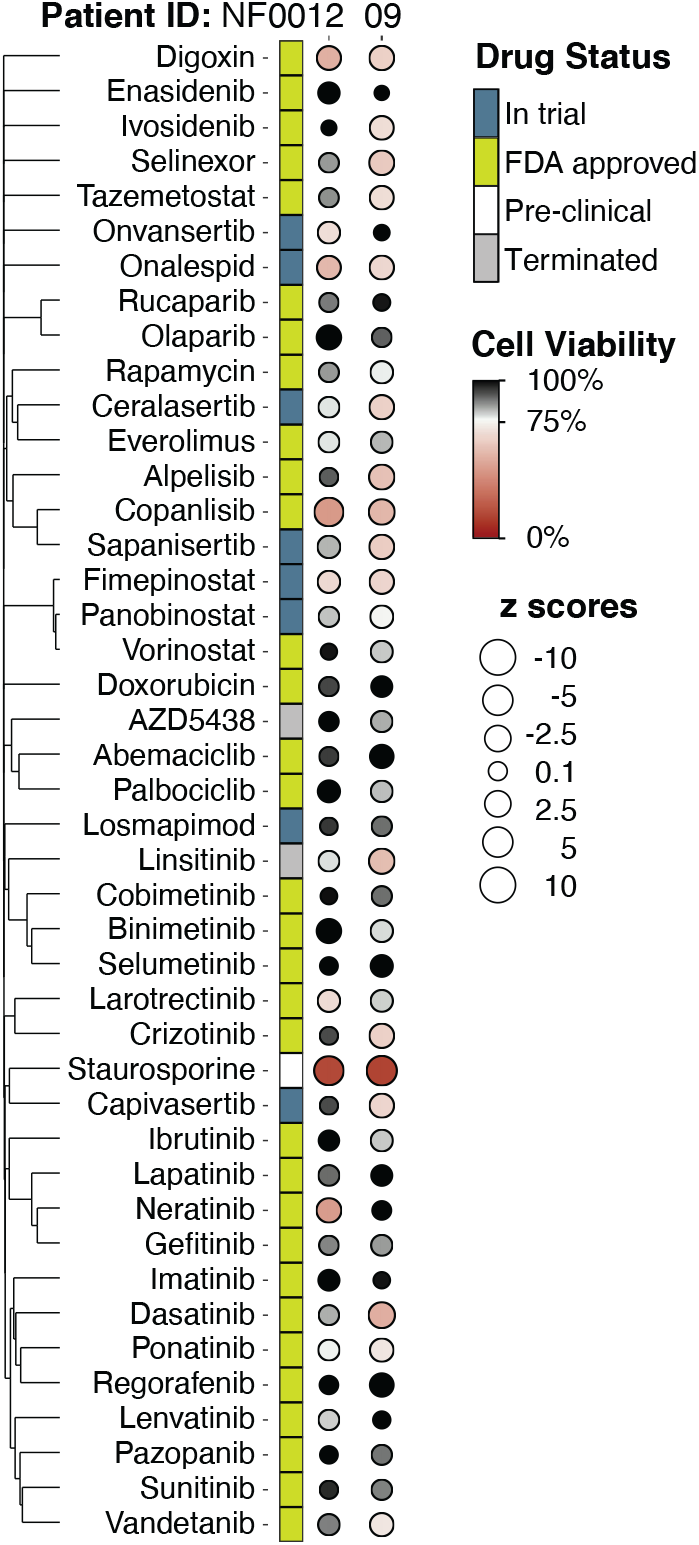
Patient-derived cNF organoids can be screened using the mini-ring organoid platform. Benign cNF tumor organoids from NF0009 and NF0012 patients were screened for response to a panel of n=43 drugs. Cell viability is normalized to vehicle (DMSO), drug approval status is included.

## Discussion

Cutaneous neurofibromas are a hallmark of neurofibromatosis type 1 syndrome and a highly penetrant feature, affecting over 90% of patients^7^. Current treatment of cNFs is limited to surgery or laser desiccation^8^. There is no systemic approach to prevent the formation of cNFs, slow down their growth or possibly shrink existing lesions, and a lack of clinically relevant, patient-derived existing model of disease to support the growth of the highly diverse cellular populations that compose cNFs.

Here we introduce a new approach toward the screening and treatment of cNFs that leverages tumor organoids. These are tractable 3D models of disease that can be established from normal tissue, benign or malignant lesions^22–24,34^. We demonstrate how viable cNF organoids generated from individual NF1 patients recapitulate the genetic variability and diverse histopathology of each individual (**Figure 1-2**).

Cutaneous neurofibromas differ from other solid tumors in their acute cell type heterogeneity. While immortalized two-dimensional models of NF1 mutant Schwann cells exist^46^, they cannot account for the cNF cellular heterogeneity and thus may not accurately recapitulate clinical responses. Our paradigm allows for the rapid establishment of tumor organoids and averts long times in culture, maintaining diverse cell populations (**Figure 1**).

We performed a detailed molecular analysis that encompasses the transcriptome, methylome, and immunohistopathology for six cNFs from a subset of n=5 NF1 syndrome patients that carry individual as well as shared pathogenic genomic alterations (**Figure 2**). At least some of the characteristics of cNFs, including their presentation and growth pattern, can be ascribed to specific *NF1* variants^12,13^. cNF organoids can be established from tissue harboring different *NF1* alterations with invariably high success rates (**Figure 1-2**). As such, organoids can rapidly expand the repertoire of models^16–19^ to investigate cNF genotype-phenotype relationships.

As media can greatly affect different cell types in the way they grow and expand in culture^30,31,34^, we screened a panel of growth conditions to determine the optimal one that preserves cell heterogeneity while also recapitulating molecular features of the parental cNF (**Figure 3-4**). Not only different cell types crucial to the establishment of cNFs are maintained (**Figure 1-4, Supplementary Figure 2**), but so is expression of extracellular matrix proteins which constitute much of the dry mass of cutaneous neurofibromas (**Figure 3d**).

The mini-ring organoid platform has been utilized to screen aggressive^23,47^ or indolent^24^ malignant tumors so far, with results available within a short timeframe from tissue procurement (5-7 days). Here, we apply this technology for the first time to the screening of heterogenous, non-malignant tumors. We can leverage this system to screen several tens (**Figure 5**) to hundreds of compounds^23,24^ and interrogate a broad range of biological pathways^24^. Given the heterogenous cellular composition of cNFs, it will be important to incorporate assays that distinguish cell types so to determine the effects that drugs have on each distinct population^45,48^.

The drug screening results suggest that there are distinct differences in sensitivity to targeted molecules that are patient-specific, possibly linked to different *NF1* alterations or additional pathogenic mutations present (**Figure 2 and 5**). Ultimately, the conditions we determined in this study lay the groundwork for rapidly testing cNF organoids established from a large cohort of NF1 patients, investigate their biology, and identify vulnerable pathways linked to specific genomic alterations.

## Acknowledgments

This work was supported by a Neurofibromatosis Therapeutic Acceleration Program grant to AS. PNNL is operated for the DOE by Battelle Memorial Institute under Contract DE-AC05-76RL01830. We thank the UCLA Translational Pathology Core Laboratory (TPCL) and the Technology Center for Genomics & Bioinformatics (TCGB) for assistance with histology and sequencing.

## Methods

### cNF tissue procurement and processing

Consented NF1 patients undergoing surgical removal of cNFs as part of their care were enrolled in this study (UCLA IRB 18-000123). Primary cNF tumors were cut into small fragments of 1-3 mm^3^ and dissociated to single cells by adding Collagenase IV (200U/ml) and incubating at 37°C with 5% CO_2_. Cells were collected every 2 hours, filtered through 70 μm cell strainer then either seeded in ring format or cryopreserved for downstream analyses.

### Media preparation

Three different base media were used: Mammocult (StemCell Technologies # 05620), DMEM/F12 (Thermo Fisher Scientific #11320033) with 10% fetal bovine serum (Life Technologies #10082-147) and StemPro™-34 SFM (Thermo Fisher Scientific # 10639011). Each medium was used as blank medium, medium with cytokines, medium with cytokines and forskolin. Cytokines including: 1X N-2 (Thermo Fisher Scientific #17502048), 10 ng/ml Neuregulin, 100 ng/ ml rhSCF, 50ng/ml rhIL-6, 1 ng/ml rhIL-3. Forskolin was used at a final concentration of 2 *µ*M.

### cNF organoids establishment

Single cell suspensions were plated around the rim of the well in the 3:4 mixture of Mammocult and Matrigel (BD Bioscience CB-40324). We seed 5’000 cells/wells for mini-rings in white 96-well plates (Corning #3603) or 100’000 cells/well for maxi-rings in 24-well plates (Corning #3527). Plates are incubated at 37 °C, 5% CO_2_ for 30 min to solidify the gel before addition of 100 *µ*l of pre-warmed medium to each well for mini-rings or 1ml of pre-warmed medium for maxi-rings. Day 3 and 4 after seeding, medium is fully removed and replaced with new pre-warmed medium. Plates are imaged daily using a Celigo S Imaging Cell Cytometer (Nexcelom) in brightfield mode to monitor daily organoid establishment and growth. For a step-by-step protocol of these procedures, see Nguyen and Soragni, STAR Protocols 2020, 1(2), 100056^25^.

### cNF organoid media screening

Five days after seeding mini-rings, the media is completely removed and wells are washed with 100 *µ*l of prewarmed PBS. We then add 50 *µ*l of 5 mg/mL dispase (Life Technologies #17105-041) to each well. After incubating at 37 °C for 25 min, plates are shaken at 80 rpm for 5 min. 75 *µ*l of CellTiter-Glo 3D Reagent (Promega #G968B) is added to each well followed by a 5-minute shake. After a 20 min incubation at room temperature and final 5-minute shake, luminescence is measured with a SpectraMax iD3 (Molecular Devices) over 500 ms of integration time. Data is normalized to base Mammocult medium and expressed as percentages^23^. Figures are generated with Prism 9 (GraphPad).

### cNF high-throughput drug screening

For drug screenings, we followed the same protocol we published^23–25^. Briefly, cells are incubated for 3 days for fresh samples, 4 days for frozen ones, to allow organoid establishment. Drug treatment is performed by complete media exchange and repeated after 24 hours. A final ATP release assay is performed 24h after the last drug treatment as described above (see cNF organoid media screening). Viability values are normalized to vehicle (DMSO) and expressed as percentages. Dot maps generated using the ggplot2 package show the 1 *µ*M drug concentration and Z scores are [(viability of drug X – viability of vehicle)/SD of vehicle]^24^.

### Sample preparation for histologic analysis

On day 5 after seeding, maxi-rings are washed with 1 ml of pre-warmed PBS, and then fixed in 500μL of 10% buffered formalin solution (VWR #89370-094). Organoids are then transferred to a 15 ml falcon tube, spun at 2000g for 5 mins followed by two washes and final removal of all the supernatant. We then add 4 μL of Histogel to the organoids pellet, which solidified on ice. The mixture of cell and histogel is placed in a cassette and followed by standard embedding, sectioning and staining^23–25^.

### Immunohistochemistry (IHC)

Slides are baked to de-paraffinize at 45°C for 20 minutes and are rehydrated in three washes of histological grade xylene (Fisher Scientific #X3S-4) followed by two washes each of ethanol (100%, 95%, and 70%) and deionized water for 10 minutes. To block naturally occurring peroxidases, Peroxidazed-1 (Biocare Medical #PX968M) is applied at room temperature for 5 minutes. Antigen retrieval is conducted using the 2100 Antigen Retriever (Atom Biologics #R2100-US) for 30 minutes at 121°C using Diva Decloaker solution (Biocare Medical #DV2004LX) for CD34 (Santa Cruz Biotechnology SC-74499), CD56 (Biocare Medical #CM164A), CD117 (Biocare Medical # CME296AK), NGFR (Biocare Medical #ACI369A), Pan Cytokeratin [AE1/AE3] (Biocare Medical #CM011A), SOX2 (Biocare Medical #ACI3109A), SOX10 (Biocare Medical #ACI3099A), and S100 (Biocare Medical #ACI3237A) antibody stainings. After slides are left to cool for 2 hours, blocking is performed at room temperature for 30 minutes with Background Punisher (Biocare Medical #BP947H). Primary antibodies are diluted in Da Vinci Green buffer (Biocare Medical #PD900L), Van Goh Yellow buffer (Biocare Medical # PD902H), or Renoir Red buffer (Biocare Medical # PD904H) at a dilution specified in the protocol chart below. 100 uL of diluted antibody is placed on the slide for primary staining, and incubation period times are listed in the protocol chart below. After 3 washes of TSBT, a MACH 3 Rabbit HRP Polymer Detection kit (Biocare Medical # M3R531H) or a MACH 3 Mouse HRP Polymer Detection kit (Biocare Medical # M3M530H) is used followed by development in Betazoid DAB kit (Biocare Medical #BDB2004).

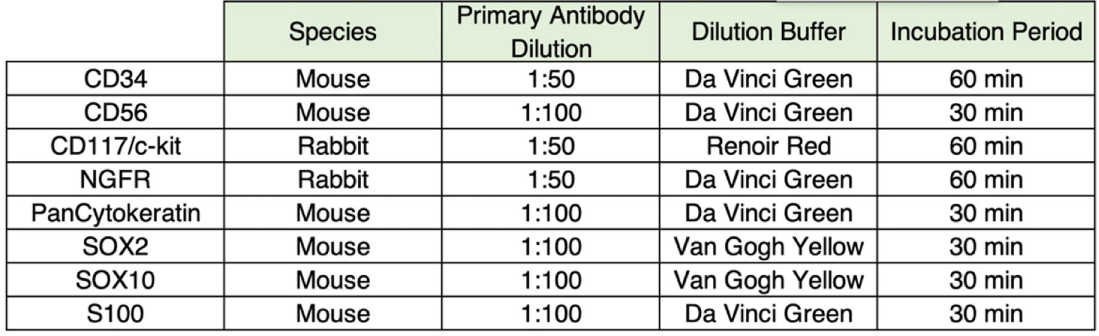

Hematoxylin (Thermo Scientific #7221) is used for counterstain. Slides are then dehydrated by two washes each of ethanol (70%, 95%, and 100%) and three washes of histological grade xylene. The slides are mounted using Permount (Fisher Scientific #SP15-100) and imaged using a Revolve Microscope (Echo Laboratories). Figures are assembled in Illustrator (Adobe).

### Whole Genome Sequencing Data Analysis

Reads were aligned to hg38 using BWA-MEM2 v2.1^49^ and converted to BAM format using SAMtools v1.10^50^. The aligned BAM files were sorted in coordinate order using Picard tools v2.23.3 SortSam function and duplicates are removed with Picard MarkDuplicates. Base quality score recalibration (BQSR) was performed with GATK v4.1.9.0 BaseRecalibrator^51^. GATK HaplotypeCaller was run on each of the split BAMs with default parameters. The VCF files generated per chromosome are merged using Picard MergeVcfs. The merged variants are then recalibrated for Indels using GATK VariantRecalibrator. Germline variants are annotated using InterVar^52^ (update-2021-08-version) to classify them according to ACMG-AMP 2015 guidelines^53^. Structural variants were called using DELLY v0.8.6^54^ on the realigned, recalibrated, reheadered BAM file for each patient. Default parameters were used except the Minimum paired-end (PE) mapping quality map-qual is set to 20. The output BCF file was then converted to VCF file using bcftools v1.11^50^. The data visualizations were created using BPG v6.0.3^55^. Basic QC metrics from the FASTQ files were generated using fastqc v0.11.9^56^. The data was aggregated from the fastqc output using the R package fastqcr v0.1.2^57^. Coverage was computed using the CollectWgsMetrics tools in Picard v2.23.3 as well as DepthOfCoverage from GATK V4.1.9.0 specifically on the *NF1* gene. The germline variants were summarized using rtg-tools v3.12^58^ on the raw VCF variants output from HaplotypeCaller.

### RNAseq and RRBS: sample preparation

On day 5 after seeding, organoids from maxi-rings are released from Matrigel by incubating at 37 °C for 20 min in 1 ml/well of 5 mg/mL dispase (Life Technologies #17105-041). Samples are spun at 1000g for 5 min, and snap frozen after removing all supernatant. We performed RNA-seq on primary tumor samples together with organoids grown in Mammocult, StemPro, and DMEM to measure the transcriptional changes that occur in the various environments. RNA was extracted from 100K cells after a 6-day growth period as organoids according to our established protocol^45^. Index-tagged sequencing libraries were prepared and sequenced by the UCLA Technology Center for Genomics & Bioinformatics (TCGB) Facility using a NovaSeq 6000 S4. For RNAseq, six samples failed quality control at the library preparation stage and are therefore not included in our analyses. These are: NF0009 with Mammocult and Mammocult + cytokines as well as NF0002 with Stempro + cytokines, Mammocult + cytokines/forskoline, DMEM + cytokines and DMEM + cytokines/forskoline.

### RNAseq and RRBS: data analysis

Raw FASTQ files were analyzed using the Salmon v1.4^59^ alignment tool with the Gencode v20^60^ annotations as part of a custom-designed workflow that we have built that consumes data directly from the NF Data Portal^29^ to add to a growing list of processed datasets that will be shared timely with the research community. To assess the cell type of origin in the organoid models we used the tumor deconvolution algorithm MCP-Counter^38^. For methylome data, raw fastq files obtained by RRBS were analyzed using BSBolt^61^ mapping against the human genome version 38 (GRCh38). Differentially methylated regions were detected using metilene v0.2-8^62^. All computational tools use to generate Figures 3 and 4 for this manuscript are available on https://github.com/PNNL-CompBio/cNFOrganoidAnalysis with the RNASeq analysis leveraging the rare disease workflows at https://github.com/Sage-Bionetworks/rare-disease-workflows/tree/main/rna-seq-workflow. These tools use the Synapse client to retrieve and store data on Synapse, which serves as the backend to the NF Data Portal at http://synapse.org/cnfOrganoids.

### Flow Cytometry

For flow cytometry, cells are spun at 600g for 5 min, resuspended in 1 ml Recovery™ Cell Culture Freezing Medium (Thermo Fisher Scientific #12648010), transferred to a cryotube and frozen at -80 degree. Cells are then thawed to perform staining and flow cytometry. APC Anti-c-Kit antibody [YB5.B8] (ab95678), Recombinant PE Anti-S100A11 antibody [EPR11172] (ab211996) and FITC Anti-CD34 antibody [4H11[APG]] (ab18227) were used to detect the mast cells, Schwann cells and fibroblast, respectively. 10 *µ*l c-kit antibody was added to a 100 *µ*l cell suspension in PBS+10% FBS then incubated for 30 minutes. After the incubation, 2 *µ*l of S100 and 1 *µ*l of CD34 antibodies are added to the mixture and incubated for an additional 30 minutes. After two washes with PBS+10% FBS, cells are resuspended in 500 *µ*l PBS+10% FBS and transferred to a 5 ml Polystyrene Round-Bottom Tube (Corning #352058) for measurement and analysis.

## Supplementary Material

### Supplementary Table

**Supplementary Table 1.**
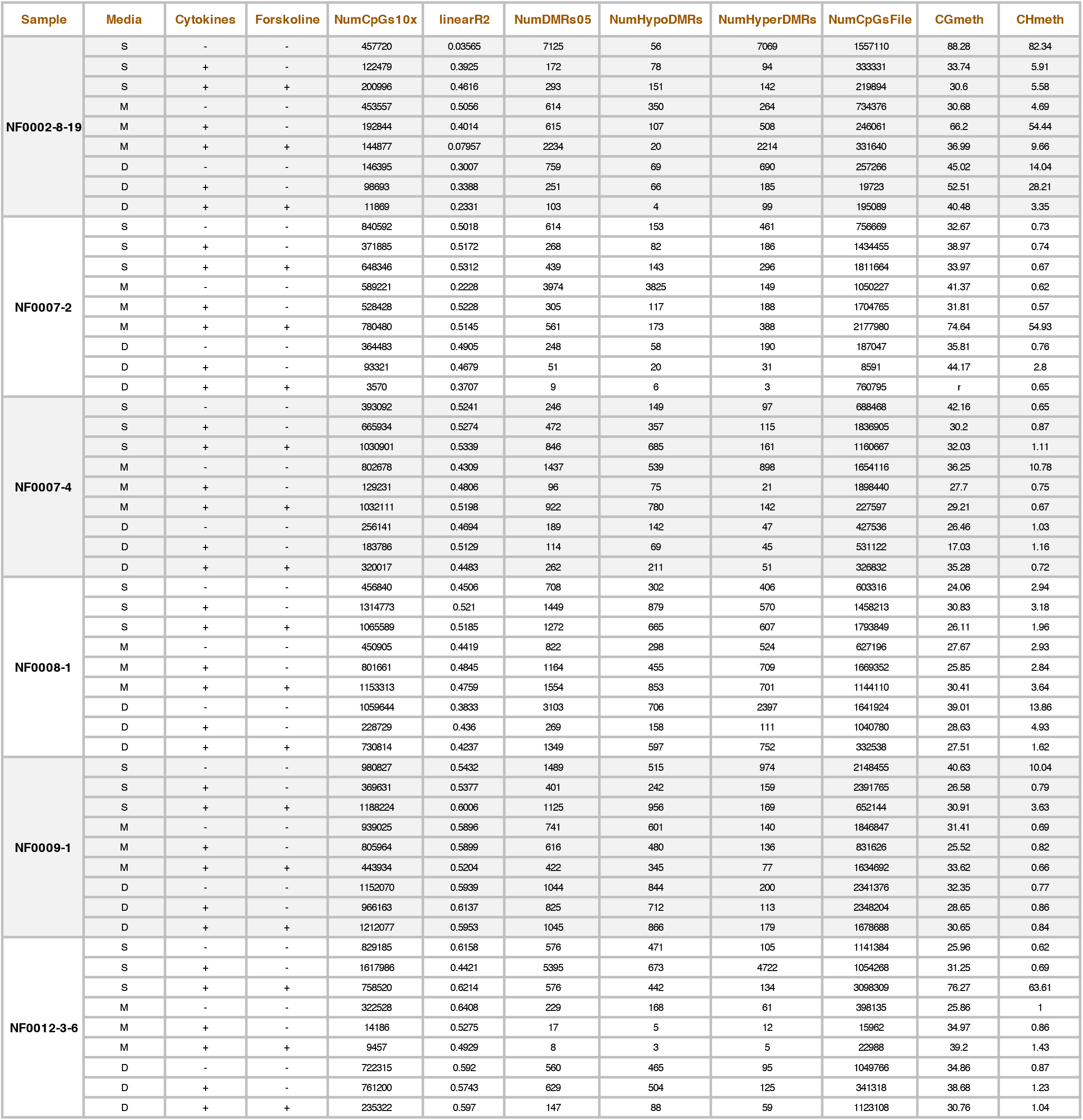
Parameters measured by RRBS analysis. Table shows methylation statistics for each treated sample against a corresponding untreated reference. NumCpGs10x corresponds to number of CpGs with coverage 10x or higher in both the treated sample and untreated control. LinearR2 corresponds to the linear r squared value for those CpGs in each treated vs. untreated sample. NumDMRs05 corresponds to the number of differentially methylated regions in treated vs untreated samples with a methylation difference larger than 50%; NumHypoDMRs are the hypomethylated and NumHyperDMRs the hypermethylated regions. NumCpGsFile are the total CpGs with coverage 10x or higher in each sample, whereas CGmeth and CHmeth corresponds to the global CG methylation and CH methylation respectively.

### Supplementary Figures

**Supplementary Figure 1.**
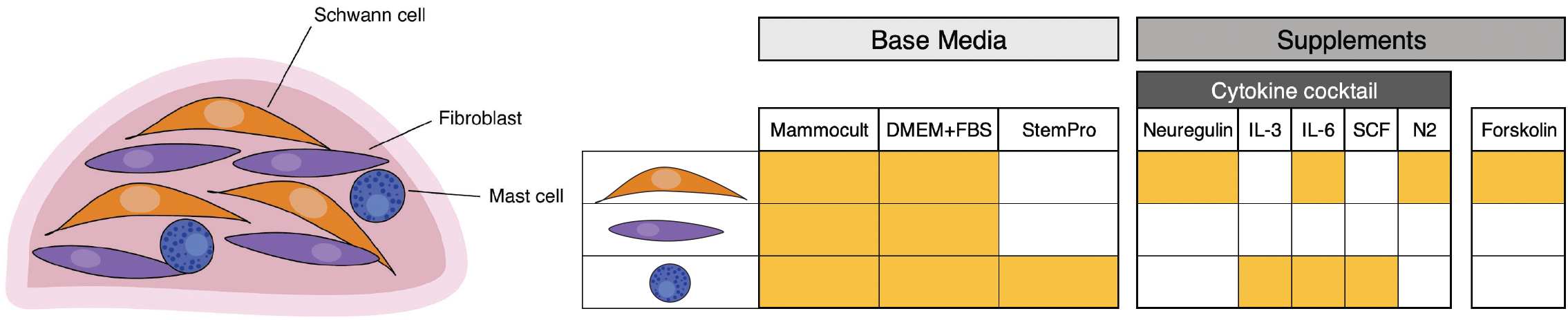
Summary of the media conditions tested. Each media or supplement was selected on the basis of literature data showing that they could support the growth of different cell types that compose cNFs.

**Supplementary Figure 2.**
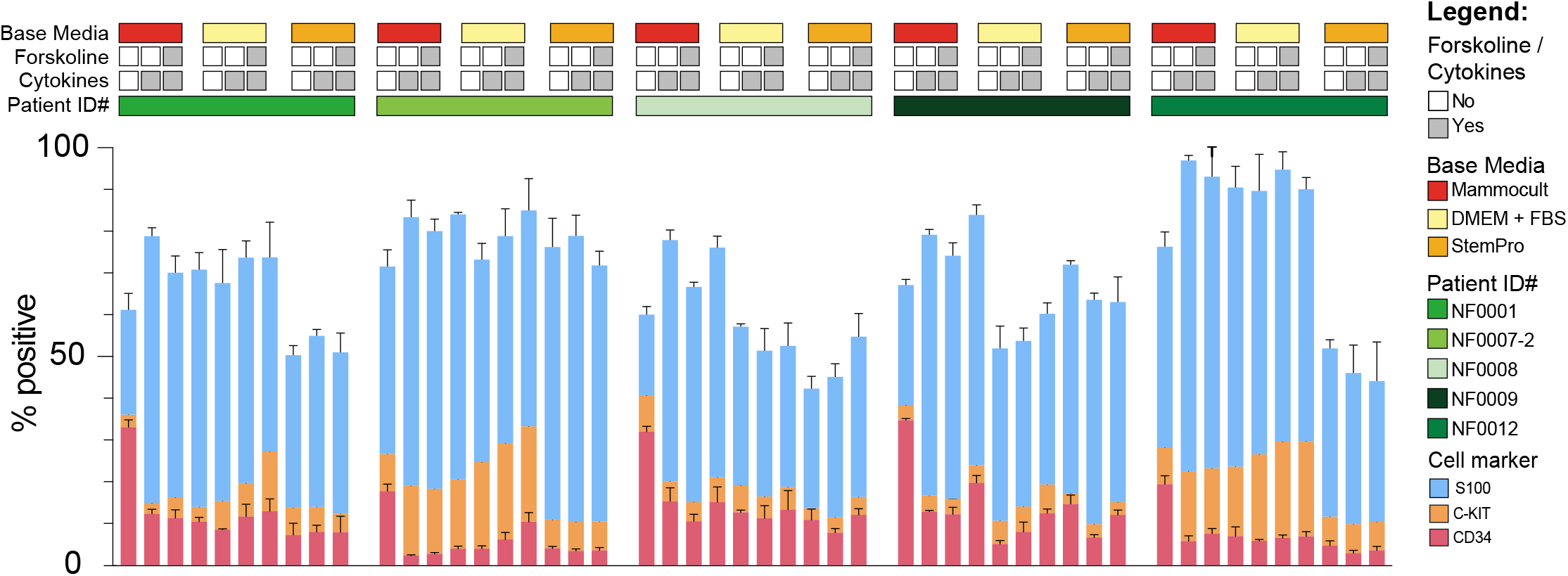
Flow cytometry of cNF tumors and organoids. The bar graphs show the percentage of cells positive for markers of Schwann cells (S100, blue), mast cells (c-kit, orange) or fibroblasts (CD34, cherry).

